# *Arid1a* protects against hepatic steatosis and insulin resistance via PPARα-mediated fatty acid oxidation

**DOI:** 10.1101/507517

**Authors:** Yu-Lan Qu, Chuan-Huai Deng, Qing Luo, Xue-Ying Shang, Jiao-Xiang Wu, Yi Shi, Lan-Wang, Ze-Guang Han

**Affiliations:** Key Laboratory of Systems Biomedicine (Ministry of Education), Shanghai Centre for Systems Biomedicine, Shanghai Jiao Tong University, Shanghai, 200240, China

**Keywords:** NAFLD, NASH, fatty acid oxidation, PPARα

## Abstract

Non-alcoholic fatty liver disease (NAFLD) and steatohepatitis (NASH) have become a worldwide health concern because of lifestyle changes, but it is still lack of specific therapeutic strategies as the underlying molecular mechanisms remain poorly understood. Our previous study indicated that deficiency of *Arid1a*, a key component of SWI/SNF chromatin remodeling complex, initiated mouse steatohepatitis, implying that *Arid1a* might be essentially required for the integrity of hepatic lipid metabolism. However, the exact mechanisms of the pathological process due to *Arid1a* loss are unclear. In the present work, we show that hepatocyte-specific deletion of *Arid1a* significantly increases susceptibility to develop hepatic steatosis and insulin resistance in mice fed with high-fat diet (HFD), along with the aggravated inflammatory responses marked by increment of serum alanine amino transferase (AST), aspartate amino transferase (AST) and TNFα. Mechanistically, *Arid1a* deficiency leads to the reduction of chromatin modification characteristic of transcriptional activation on multiple metabolic genes, especially *Cpt1a a*nd *Acox1*, two rate-limiting enzyme genes for fatty acid oxidation. Furthermore, our data indicated that *Arid1a* loss promotes hepatic steatosis by downregulating PPARα, thereby impairing fatty acid oxidation which leads to lipid accumulation and insulin resistance. These findings reveal that targeting ARID1a might be a promising therapeutic strategy for NAFLD, NASH and insulin resistance.

## Introduction

Non-alcoholic fatty liver disease (NAFLD), as characterized by hepatic steatosis, insulin resistance and chronic low-grade inflammation, is the leading cause of chronic liver disease, and may process to non-alcoholic steatohepatitis (NASH), cirrhosis and hepatocellular carcinoma (1,2). Ectopic lipid accumulation is proposed to be the root cause of NAFLD, leading to hepatic steatosis and insulin resistance (3). Fatty acid oxidation (FAO) is a multistep process that involves the degradation of fatty acids (FAs) by the sequential removal of two-carbon units from the acyl chain to produce acetyl-CoA, and there are compelling evidences show that undermined hepatic FAO is one of the central processes in hepatic lipid disorders (4). Inadequate oxidation of fatty acid would result in aberrant lipid accumulation and massive steatosis (2). However, the underlying mechanisms for this process are complex, multifactorial and heterogeneous, which hinder the exploration of exact pathological processes and therapeutic methods.

SWI/SNF chromatin remodeling complex manipulates the accessibility of regulatory transcriptional factors and thereby controls the transcriptional activation and repression of some genes related to physiological functions (5,6). So far, the effects of SWI/SNF complex members on nutrient metabolism remains less clear. It was reported that hepatocyte-specific deletion of SNF5, another subunit of SWI/SNF complex, caused glycogen storage deficiency and energetic metabolism impairment (7). Baf60a and Baf60c regulated metabolic gene programs in the liver and skeletal muscle (8,9), and even, Baf60a controlled the transcriptional programs of some genes responsible for hepatic bile acid synthesis and intestinal cholesterol absorption (10). Moreover, expression of BAF60a could stimulate fatty acid β-oxidation in cultured hepatocytes and ameliorated hepatic steatosis *in vivo (11).*

ARID1a, also known as Baf250a, a subunit of SWI/SNF chromatin remodeling complex, may facilitate the access of transcription factors and regulatory proteins to genomic DNA. Loss of ARID1A may result in the structural and functional alterations of SWI/SNF complex, which leads to transcriptional dysfunction through disruption of nucleosome sliding activity, assembly of variant SWI/SNF complexes, targeting to specific genomic loci, and recruitment of coactivator/corepressor activities (12-17). Our group previously demonstrated hepatocyte-specific *Arid1a* deficiency initiated mouse hepatic steatosis and steatohepatitis (18), suggesting that *Arid1a* might participate in lipid mechanism, however, the exact pathological process and the underlying molecular mechanism were unclear. In this study, we employed hepatocyte-specific *Arid1a* knockout mice administrated with high fat diet (HFD), and proved that wild type *Arid1a* protected mice against hepatic steatosis and insulin resistance. Mechanistically, *Arid1a* directly controlled the transcriptional activation of multiple metabolic genes by erasing H3K4me3 marks, and downregulated *PPARα* which thus inhibited fatty acid oxidation to regulate lipid metabolism and insulin sensitivity. In general, the finding that *Arid1a* may regulate lipid mechanism via directly and indirectly modulating the transcription of fatty acid oxidation-related genes provides a promising therapeutic approach for NAFLD, NASH and insulin resistance.

## Results

### *Arid1a* is required for regulating insulin sensitivity and glucose tolerance

Insulin resistance is a vital part in the development of hepatic steatosis (19). To directly address the effect of *Arid1a* deletion on glucose homeostasis and insulin sensitivity, liver-specific *Arid1a* knockout (*Arid1a*^*LKO*^) mice and *Arid1a*^*F/F*^ mice were placed on a normal chow diet (CD) or high-fat diet (HFD), and then metabolic phenotypes of these mice were characterized. Interestingly, despite of similar food intake **(Supplementary Fig. 1A)**, *Arid1a*^*LKO*^ mice gained much more body weights than *Arid1a*^*F/F*^ mice when placed on a HFD, whereas no difference was statistically identified between the *Arid1a*^*LKO*^ and *Arid1a*^*F/F*^ littermates under CD **(Fig. 1A-B)**. In parallel, compared to *Arid1a*^*F/F*^ littermates fed on HFD, *Arid1a*^*LKO*^ mice showed significantly elevated levels of fasting blood glucose **(Fig. 1C)** and insulin **(Fig. 1D)**, as well as the markedly decreased glucose tolerance via glucose tolerance test (GTT) **(Fig. 1E)** and insulin sensitivity shown by insulin tolerance test (ITT) **(Fig. 1F)**. Furthermore, insulin sensitivity was assessed by measuring the phosphorylated Protein Kinase B (pAKT) level in livers of *Arid1a*^*LKO*^ mice fed on HFD. As expected, after an injection of insulin, pAkt level was significantly decreased in the livers of *Arid1a*^*LKO*^ mice **(Fig. 1G)**, as compared to that of *Arid1a*^*F/F*^ littermates. However, no significant difference in blood glucose and insulin levels were found between *Arid1a*^*LKO*^ and *Arid1a*^*F/F*^ group fed on CD during GTT and ITT tests **(Supplementary Fig. 1B-E)**. Collectively, these data indicate that hepatocyte-specific *Arid1a* deletion damages insulin sensitivity, thereby leading to insulin resistance and glucose tolerance of *Arid1a*^*LKO*^ mice under HFD challenge.

**Figure 1.**
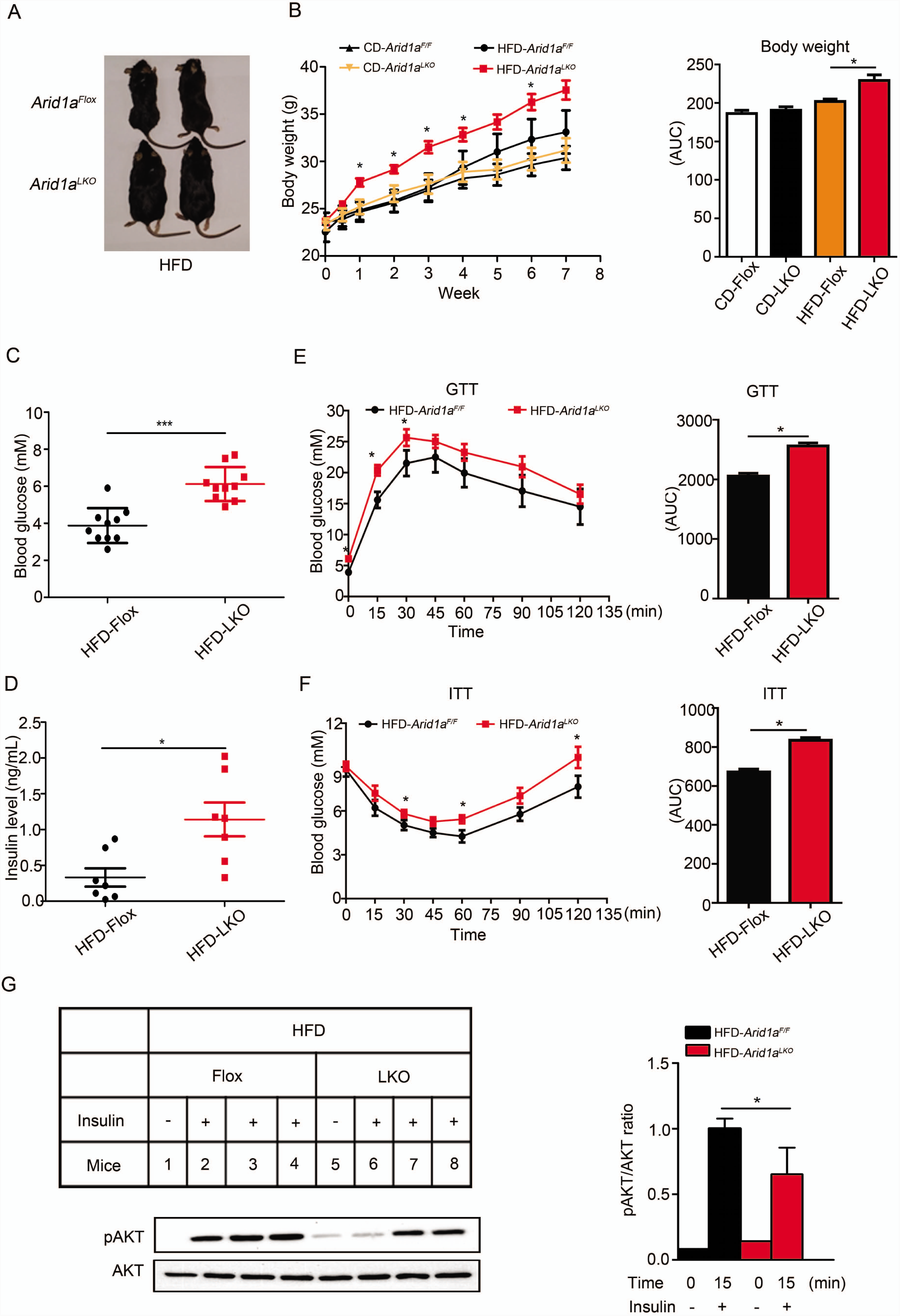
Hepatic *Arid1a* deletion alters glucose tolerance and promotes insulin resistance. (A) Representative images of *Arid1a*^*LKO*^ and *Arid1a*^*F/F*^ littermates after HFD feeding for 12 weeks. (B) Left, growth curve of *Arid1a*^*LKO*^ and *Arid1a*^*F/F*^ littermates on a CD or HFD; Right, quantitation of body weight (n = 9-10). (C) Fasting blood glucose level of mice on HFD (n = 10). (D) ELISA-determined fasting serum insulin level of mice on HFD (n = 7). (E) Left, measurement of plasma glucose during glucose tolerance test (GTT) of mice on HFD; Right, the calculated AUCs (area under curves) in *Arid1a*^*LKO*^ and *Arid1a*^*F/F*^ mice on HFD (n = 8-10). (F) Left, measurement of plasma glucose during insulin tolerance test (ITT) of mice on HFD; Right, the calculated AUCs (area under curves) in *Arid1a*^*LKO*^ and *Arid1a*^*F/F*^ mice on HFD (n = 8-10). (G) Left, total and phosphorylated AKT levels in the livers of *Arid1a*^*LKO*^ and *Arid1a*^*F/F*^ mice on HFD after intraperitoneal insulin injection; Right, the expression levels of the protein as phosphorylated- to total-protein ratio (n = 3). Statistical analysis was performed. Values are mean ± SD. * *P* < 0.05. *** *P* < 0.001.

### HFD aggravates *Arid1a* deficiency-induced hepatic steatosis, fibrosis and inflammation

To define the role of *Arid1a* on the development of non-alcoholic fatty liver, we carried out histological comparisons between the livers from 4 months-age *Arid1a*^*LKO*^ and *Arid1a*^*F/F*^ mice. Consistent with our previous data, *Arid1a*^*LKO*^ mice fed with CD exhibited hepatic dysplasia, steatosis and fibrosis, as compared to *Arid1a*^*F/F*^ mice (18). However, HFD significantly aggravated these phenotypes of *Arid1a*^*LKO*^ mice, as evidenced by quantitatively analyses on histology according to the NAFLD Activity Score (NAS) (20) which measures degree of steatosis, lobular inflammation, and hepatocyte ballooning **(Fig. 2A and B)**. In addition, more lipid droplets were accumulated in the livers of *Arid1a*^*LKO*^ mice challenged with HFD, as visualized by Oil Red O staining **(Fig. 2C)**. In parallel with these morphologic changes, the *Arid1a*^*LKO*^ mice fed on HFD exhibited worse liver function, as indicated by significant elevation of serum AST, ALT and ALP levels **(Fig. 2D-F)**. Moreover, the mRNA levels of inflammatory-related genes including *Tnfα, Arg1, Mcp1 and Cd11b* were also markedly elevated in HFD-fed *Arid1a*^*LKO*^ mice compared to those in the *Arid1a*^*F/F*^ mice **(Fig. 2G)**. Correspondingly, blood concentration of TNF-α **(Fig. 2H)** was also significantly increased in obese *Arid1a*^*LKO*^ mice, although serum level of IL-6 had no significant change **(Fig. 2I)**. Taken together, these data indicate that HFD aggravates *Arid1a* deficiency-induced hepatic steatosis, fibrosis and inflammation, suggesting that the *Arid1a* deficiency-induced dysfunction of lipid mechanism could be crucial to the development of hepatic disease.

**Figure 2.**
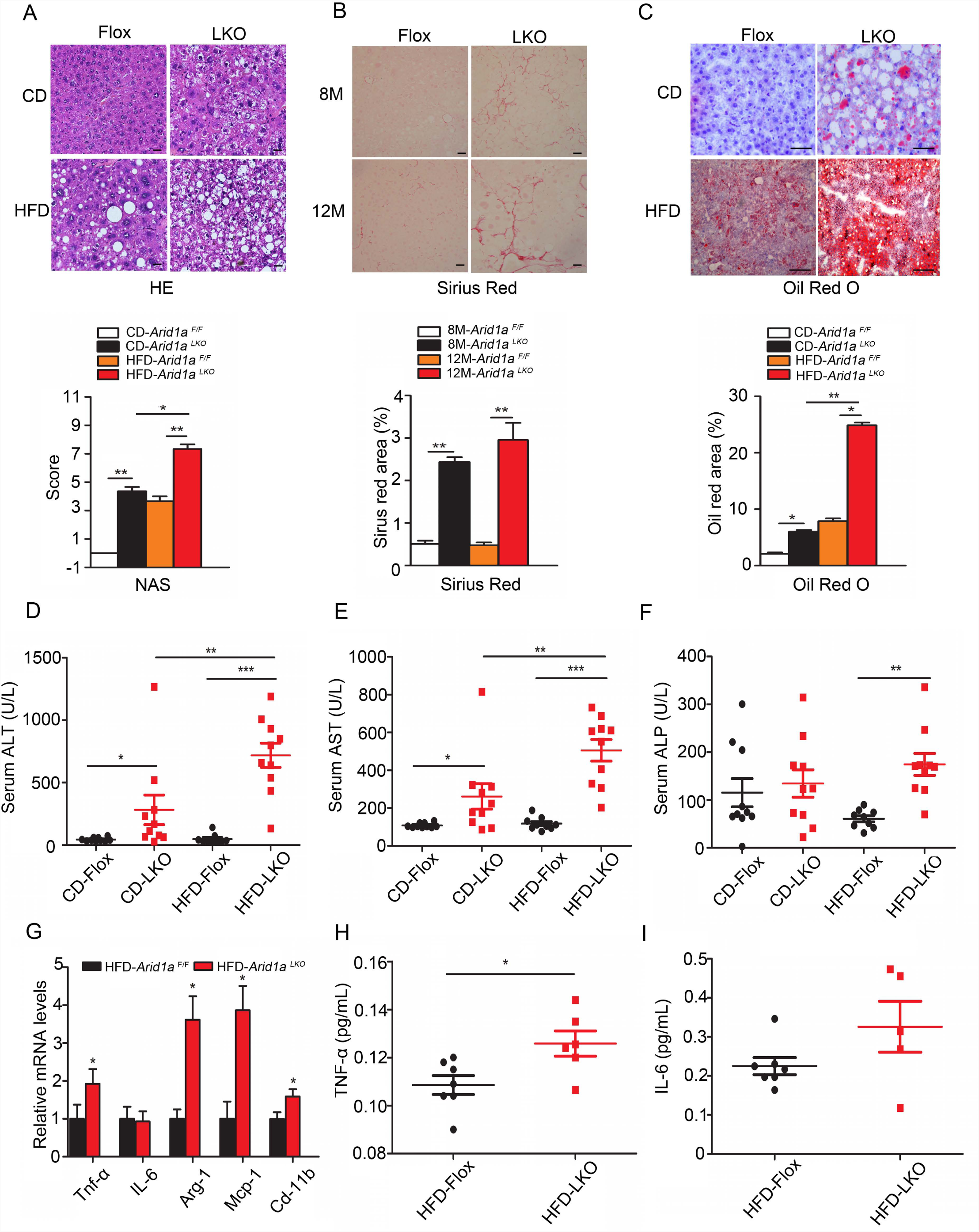
HFD aggravates *Arid1a* deficiency-induced hepatic steatosis, fibrosis and inflammation. (A) Upper, H&E-stained histological images of liver sections from CD and HFD-fed *Arid1a*^*LKO*^ and *Arid1a*^*F/F*^ mice; Down, NAS scores for *Arid1a*^*LKO*^ and *Arid1a*^*F/F*^ mice at 4 months of age. Scale bar, 50 μm. (B) Upper, representative images of Sirius red-stained liver sections from HFD-fed *Arid1a*^*LKO*^ and *Arid1a*^*F/F*^ mice; Down, fibrosis area was quantified. Scale bar, 50 μm. (C) Lipid droplets visualized by Oil Red O staining in livers; Down, lipid droplet area was quantified. Scale bar, 50 μm. (D-F) Serum levels of ALT (D), AST (E) and AFP (F) of male mice with the indicated diet and genotypes (n = 9-10). (G) qPCR analysis of inflammatory genes in livers of mice on HFD (n = 7-8). (H-I) Serum levels of TNFα (H) and IL-6 (I) in mice fed HFD for 12 weeks (n = 5-7). Statistical analysis was performed. Values are mean ± SD, * *P* < 0.05; ** *P* < 0.01; *** *P* < 0.001.

### *Arid1a* deletion leads to hepatic steatosis possibly through FAO impairment

Metabolic imbalance between lipid acquisition and removal in the liver is the first step in the pathophysiology of NAFLD (21). To explore the effects of *Arid1a* on hepatic lipid metabolism, the mice were fed with CD or HFD for 12 weeks and then sacrificed to observe their phenotypes. Significantly, the liver weights of the *Arid1a*^*LKO*^ mice fed on HFD were much higher than those of the *Arid1a*^*F/F*^ mice **(Fig. 3A)**. Additionally, both subcutaneous white adipose tissue (WAT) and inguinal WAT fat pads were larger in *Arid1a*^*LKO*^ mice **(Fig. 3B and C)**, indicative of decreased adipose tissue homeostasis. Afterwards, blood lipid testing revealed that plasma cholesterol (TCHO) as well as low-density lipoprotein (LDL) and high-density lipoprotein cholesterol (HDL) cholesterol levels, but not non-esterified fatty acids (NEFA), were much higher in *Arid1a*^*LKO*^ animals **(Fig. 3D-F)**. Surprisingly, plasma triglycerides (TG) of *Arid1a*^*LKO*^ mice was reduced **(Fig. 3D).** Furthermore, we determined liver lipids. TCHO and TG content in the livers from HFD-fed *Arid1a*^*LKO*^ mice were significantly increased, where hepatic TG in *Arid1a*^*LKO*^ mice were even 8-fold higher than those in *Arid1a*^*F/F*^ mice **(Fig. 3G)**, indicating that the hyperlipidemia of *Arid1a*^*LKO*^ mice on HFD is likely caused by hepatic *Arid1a* loss, although there is barrier to block the secretion of liver TG to blood.

**Figure 3.**
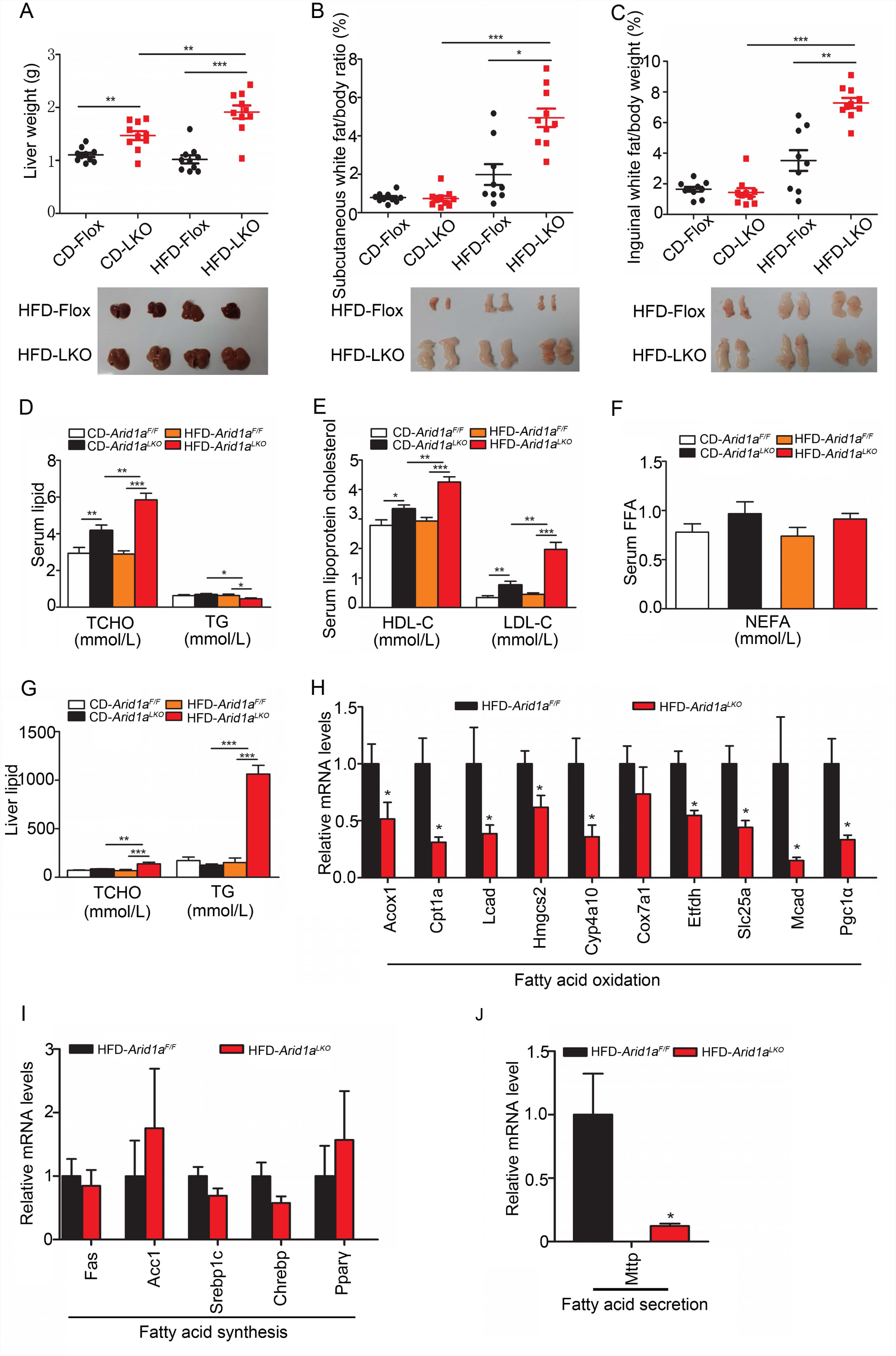
*Arid1a* deprivation causes severe lipid accumulation by impairing FAO. (A) Upper, determination of liver weights of *Arid1a*^*LKO*^ and *Arid1a*^*F/F*^ mice at 4-months age; Down, representative images of livers of *Arid1a*^*LKO*^ and *Arid1a*^*F/F*^ mice on HFD (n = 10). (B-C) Upper, determination of subcutaneous WAT (B) and inguinal WAT (C) weights of *Arid1a*^*LKO*^ and *Arid1a*^*F/F*^ mice; Down, representative images of subcutaneous WAT and inguinal WAT of mice on HFD (n = 9-10). (D-F) TG, TCHO (D), lipoprotein-cholesterol (E) and NEFA (F) concentrations in the serum of *Arid1a*^*LKO*^ and *Arid1a*^*F/F*^ mice (n = 7-10). (G) Analysis of TG and TCHO contained in the livers (n = 7-10). (H-J) Real-time analysis of genes involved in fatty acid oxidation (H), synthesis (I) and secretion (J) processes (n = 7). Statistical analysis was performed. Values are mean ± SD, * *P* < 0.05; ** *P* < 0.01; *** *P* < 0.001.

Based on the observations that dysregulated lipid metabolism was induced by *Arid1a* deletion, we assessed transcriptional expression of genes related to lipid acquisition and removal, including fatty acid synthesis, oxidation and TG secretion. The mRNAs encoded by these genes involved in FAO (*Pparα, Cpt1a, Acox1, Pgc1α, Hmgcs2*, etc.) and secretion (*Mttp*) were significantly reduced in the *Arid1a*^*LKO*^ mice with HFD administration whereas there was no obvious difference of genes expression related to fatty acid synthesis (*Fas, Acc1, Pparγ, Sreb1c* and *Chrebp*) between *Arid1a*^*LKO*^ and *Arid1a*^*F/F*^ mice **(Fig. 3H-J)**, suggesting that the reduced FAO and impaired TG secretion may contribute to hepatic lipid accumulation in *Arid1a*^*LKO*^ mice.

### *Arid1a* deficiency augments FFA-induced lipid accumulation and insulin resistance

To directly explore the role of *Arid1a* in FAO process, we investigated whether *Arid1a* can regulate free fatty acids (FFA)-induced lipid accumulation and insulin signaling in the hepatocytes isolated from *Arid1a*^*F/F*^ mice, where *Arid1a* was deleted by adenovirus containing *Cre* (Ad-Cre) infection **(Fig. 4A).** Interestingly, *Arid1a* deficiency significantly augmented the oleic acid (OA, the unsaturated fatty acid) and palmitic acid (PA, the saturated fatty acid) -induced lipid accumulation in these hepatocytes **(Fig. 4B)**, indicating that *Arid1a* was required for FAO process to break down these saturated and unsaturated fatty acid molecules. Later on, we assessed the alterations of insulin response in the primary hepatocytes with *Arid1a* loss. The results showed that, regardless of the presence or absence of OA, *Arid1a* deficiency led to decreased phosphorylation levels of AKT and GSK-3β in hepatocytes upon insulin stimulation **(Fig. 4C).** Similar results were observed in hepatocytes isolated from tamoxifen-induced *Arid1a* knockout mice, as well as in hepatocellular carcinoma (HCC) cell lines MHCC-97H and SK-Hep1-6 with *Arid1a* knockout by CRISPR-Cas9 system upon insulin treatment **(Supplementary Fig. 2)**.

**Figure 4.**
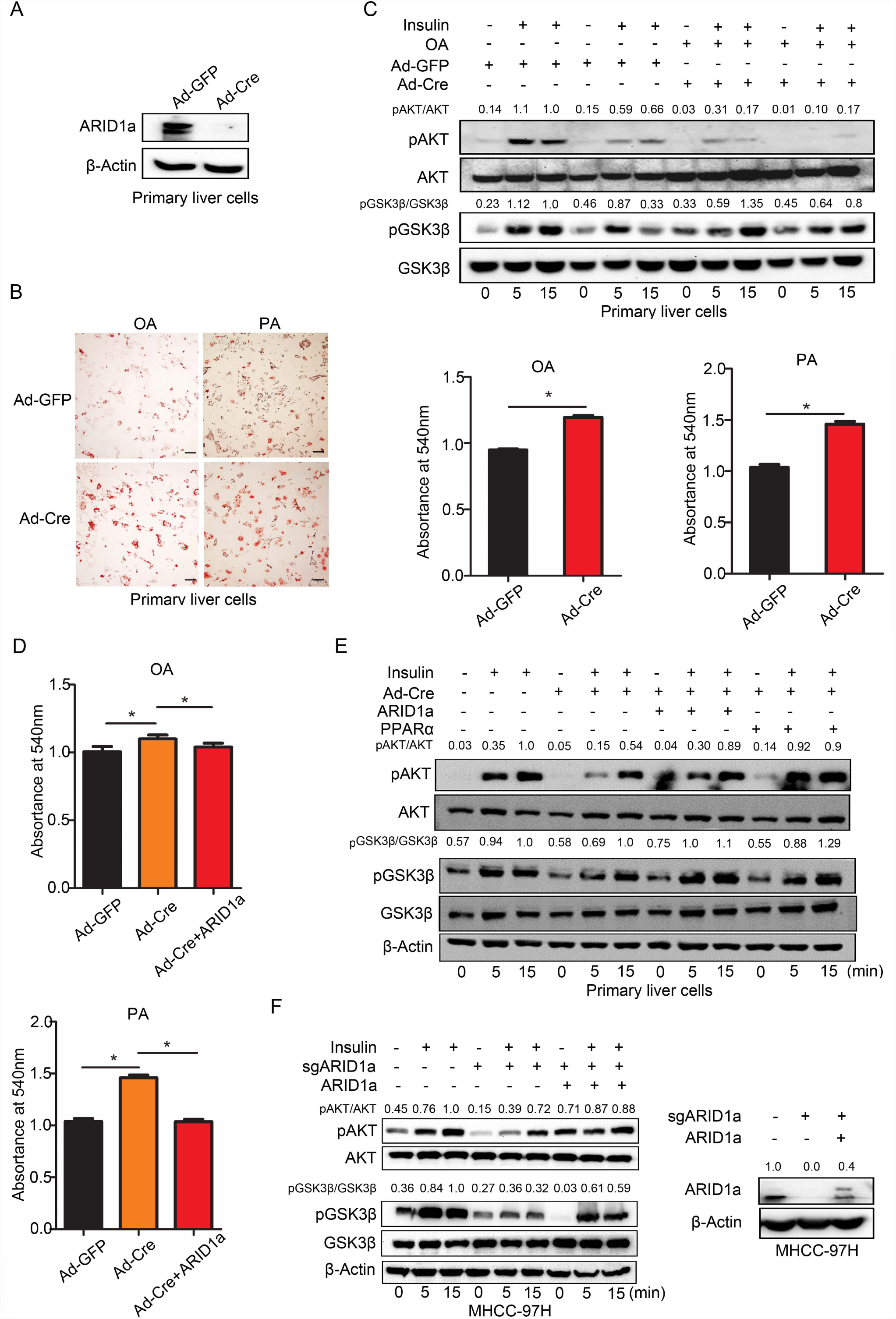
*Arid1a* deficiency attenuates FAO and insulin sensitivity in hepatocytes. (A) Western blotting confirmed that *Arid1a* was deleted in hepatocytes infected with Ad-Cre. (B) Left, representative images of Oil Red O staining in Ad-GFP or Ad-Cre transfected hepatocytes with oleic acid (OA) or palmitic acid (PA) stimulation; Right, lipid content analysis. (C) Phosphorylation levels of AKT at serine 473 and GSK-3β at serine 9 were assessed in hepatocytes with or without OA stimulation. (D) Ectopic ARID1a expression reverses lipid accumulation in *Arid1a*^*-/-*^ hepatocytes. (E-F) ARID1a restoration reverses insulin resistance in hepatocytes (E) and MHCC-97H cell lines (F) with *Arid1a* deletion. Statistical analysis was performed. Values are mean ± SD, * *P* < 0.05.

Next, we performed the rescue assay on the *Arid1a*^*-/-*^ primary mouse hepatocytes via ectopic ARID1a expression. Remarkably, the rescued ARID1a expression resulted in the restoration of lipid clearance in *Arid1a*-deficient hepatocytes under both OA and PA challenge, which was evidenced by quantitative analysis of Oil Red O staining **(Fig. 4D and 6F)**. Furthermore, the phosphorylation levels of AKT and GSK3β were also significantly recovered by ectopic ARID1a expression, and similar results were acquired in the MHCC-97H cells with *Arid1a* deletion **(Fig. 4E and F).** Altogether, these data indicate that *Arid1a* may disrupt the fatty acid β-oxidation as a critical process for lipid metabolism regulation.

### *Arid1a* loss down-regulates essential genes in FAO and Ppar signaling pathways

Then we performed RNA sequencing on the mouse hepatocytes infected with Ad-Cre or Ad-GFP to figure out the *Arid1a*-regulated downstream genes (GEO). Significantly, pathway analysis using the GO and KEGG databases revealed that the down-regulated genes induced by *Arid1a* deficiency were enriched in lipid and fatty acid metabolic processes, PPAR and insulin signaling pathways, respectively **(Fig. 5A and B)**. Additionally, we performed the gene set enrichment analysis and found that fatty acid metabolism was the Top5 ranked among the ‘Hallmark’ gene sets **(Fig. 5C)**. Many genes related to fatty acid metabolism and PPAR signaling pathways were downregulated in the *Arid1a* deficient group **(Figure 5D)**, some of which were consistent with qRT-PCR results in the mouse liver (**Fig. 3H**), suggesting that *Arid1a* loss induces the dysregulated lipid metabolism and PPAR function.

**Figure 5.**
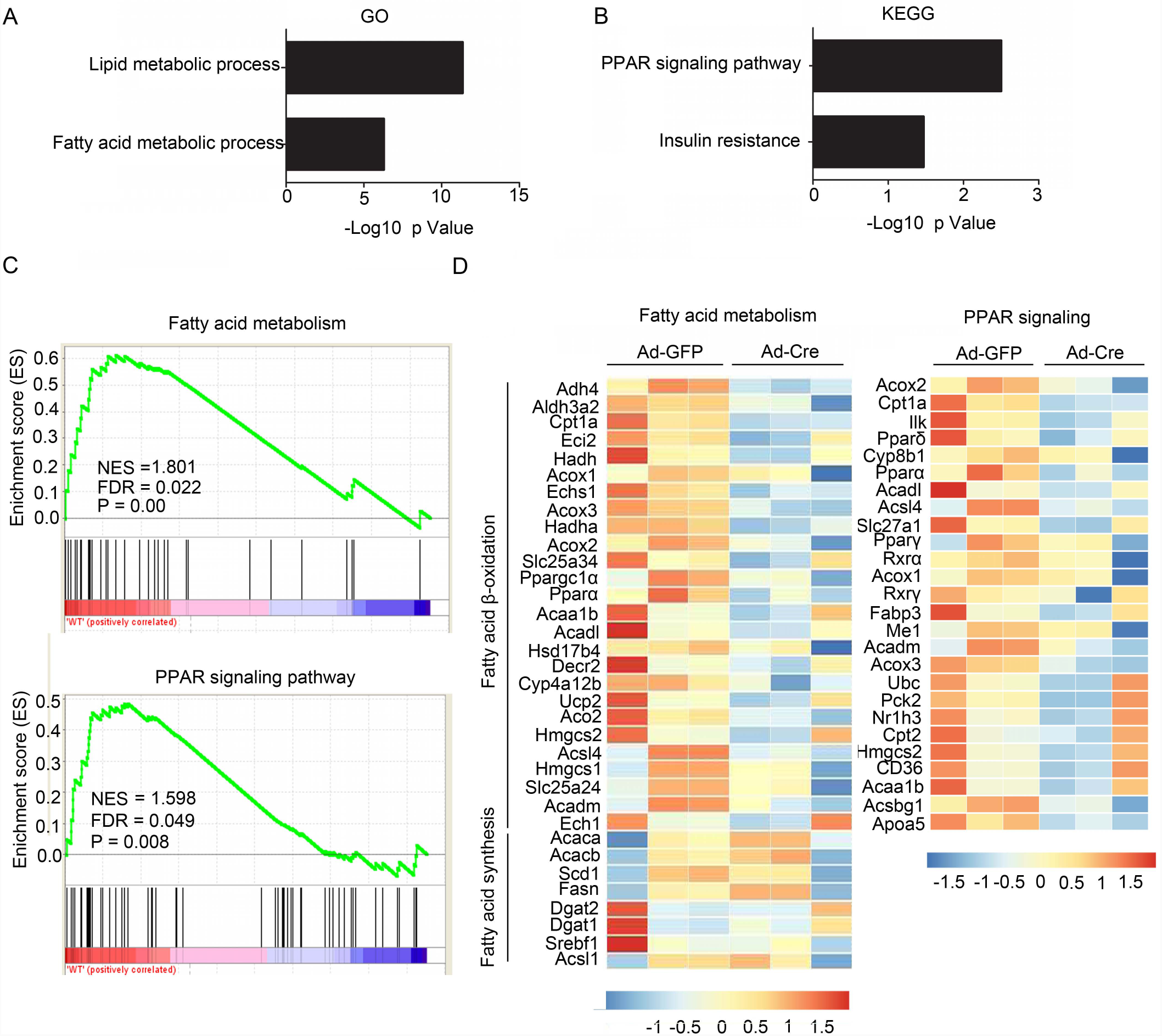
*Arid1a* deprivation causes defects in lipid metabolic process and PPAR signaling. (A-B) GO (A) and KEGG (B) pathway analysis of the down-regulated genes based on RNA-seq data of *Arid1a*-deficient hepatocytes. (C) GSEA showed downregulation of the genes related to fatty acid metabolism and PPAR signaling pathways in *Arid1a*-deficient hepatocytes. (D) Heat map of mRNA levels of the genes involved in fatty acid metabolism and PPAR signaling. Red and blue depict higher and lower gene expression, respectively. Color intensity indicates magnitude of expression differences (n = 3).

We further validated some RNA-seq revealed differentially expressed genes in hepatocytes by qRT-PCR, showing that *Arid1a* deficiency caused a significant reduction of FAO genes, such as *Pparα, Acox1, Cpt1α*, and *Hmgcs2*, in the primary mouse hepatocytes (**Fig. 6A)** and immortalized hepatocytes with SV40 expression **(Supplementary Fig. 3)**. In contrast, the mRNA levels of lipogenic genes, including *Acc1, Chrebp, Srebp1c* and *Fas*, were not obviously changed in hepatocytes with *Arid1a* deletion. Moreover, the mRNA expression levels of key gluconeogenic enzymes were determined. As shown in Figure 6A, glucose 6-phosphatase (*G6pase*), but not phosphoenolpyruvate carboxykinase (*Pepck*) mRNAs were increased 2.5-fold in the hepatocytes with *Arid1a* deficiency **(Fig. 6A)**. qRT-PCR and western blotting assay also revealed that insulin receptor substrate 1 (*Irs1*) was significantly down-regulated in mouse hepatocytes as well as in MHCC-97H cells with *Arid1a* knockout **(Supplementary information, Fig. 4)**. Consistently, ectopic ARID1a expression recovered the expression of *Pparα, Acox1, Cpt1a, Hmgcs2*, and *Pgc1α* in the mouse hepatocytes with *Arid1a* deletion **(Fig. 6B)**.

**Figure 6.**
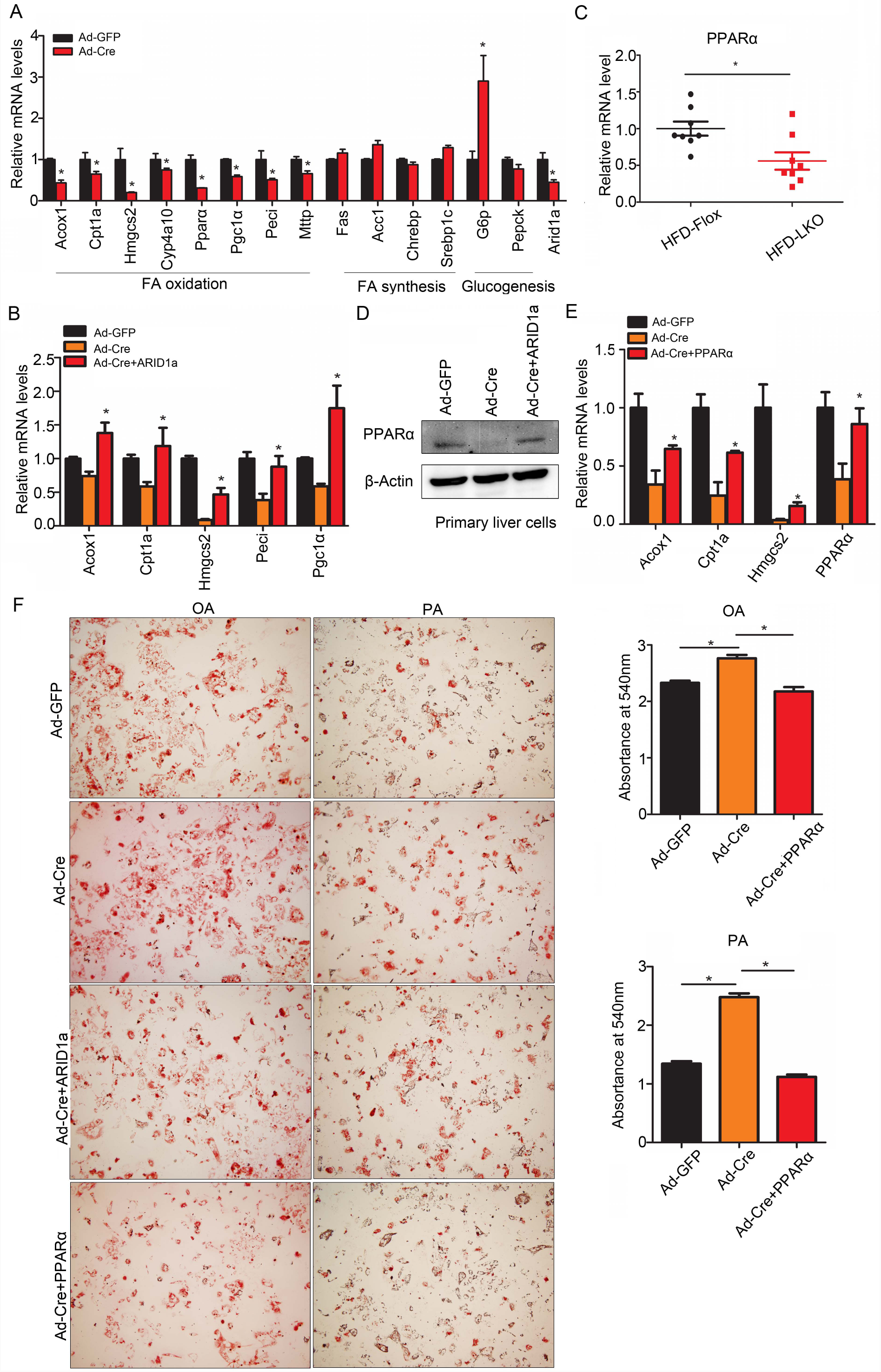
Ectopic PPARα expression protects against FAO impairment and lipid accumulation in hepatocytes with *Arid1a* deletion. (A) Real-time analysis of the genes expression involved in lipid or glucose metabolism in hepatocytes transfected with Ad-GFP or Ad-Cre. (B) Ectopic ARID1a expression stimulates mRNA levels of fatty acid oxidation-related genes in *Arid1a*^*-/-*^ hepatocytes. (C) Real-time analysis of PPARα expression in mouse livers (n = 8). (D) PPARα expression was restored by ARID1a overexpression in *Arid1a*^*-/-*^ hepatocytes. (E) *Arid1a* deletion-induced *Cpt1a* and *Acox1* downregulation was partially abrogated by PPARα overexpression in hepatocytes. (F) Left, lipid droplets visualized by Oil red O staining in hepatocytes; Right, quantitation of lipid content in hepatocytes. Statistical analysis was performed. Values are mean ± S D, * *P* < 0.05.

Among the mentioned effector genes of ARID1a, we paid particularly attention to Pparα, the major regulator of fatty acid oxidation (22), which was largely reduced in the liver and hepatocytes without *Arid1a* **(Fig. 6A and C)**. Remarkably, western blotting assay confirmed that Pparα was recovered by ectopic ARID1a expression in the *Arid1a*^*-/-*^ hepatocytes **(Fig. 6D)**, suggesting that the down-regulated Pparα might be critical for *Arid1a* deletion-induced lipid accumulation and insulin resistance. Later on, we investigated whether induction of PPARα can improve FAO in *Arid1a*^*-/-*^ hepatocytes. Interestingly, the results showed that the enforced expression of PPARα partially recovered the defective expression of its targeting downstream genes *Acox1, Cpt1a*, and *Hmgcs2* in *Arid1a*^*-/-*^ hepatocytes **(Fig. 6E)**. Consequently, ectopic PPARα expression alleviated the lipid accumulations induced by OA and PA **(Fig. 6F**), and insulin sensitivity, indicated by pAKT and pGSK3β in mouse *Arid1a*^*-/-*^ hepatocytes **(Fig. 4E)**. The results testified that *Arid1a* deficiency-induced transcription repression of Pparα and related fatty acid oxidation genes could be responsible for hepatic lipid accumulation and steatosis.

### *Arid1a* deficiency decreases H3K4me3 and chromatin accessibility on promoters of metabolic genes

It is noteworthy that Arid11a may facilitate access of transcription factors and regulatory proteins to the genomic DNA, and thus regulates the transcription of downstream genes. Therefore, we proposed that Arid1a may impact on the local epigenetic landscape to regulate transcription activity of key metabolic genes. Here, we firstly analyzed the published ChIP-seq dataset (GSE65167) (23). Bioinformatic analysis based on GO and KEGG database indicated that the *Arid1a* targeted genes are highly associated with lipid, fatty acid metabolic processes and PPAR signaling **(Fig. 7A and B)**. Then we integrated the *Arid1a*-regulated genes, indicated by RNA-seq data, in mouse hepatocytes, with the genes containing Arid1a-binding peaks on promoter, showing that two key FAO related genes *Acox1* and *Cpt1a* have the Arid1a-binding peaks **(Fig. 7C)**. Notably, Cpt1a is a rate-limiting enzyme involved in mitochondrial β-oxidation, catalyzing the esterification of long-chain acyl-CoAs to L-carnitine for their transportation into the mitochondria (24). While Acox1 is the first and a rate-limiting enzyme in the peroxisomal β-oxidation which catalyzing the desaturation of very-long-chain acyl-CoAs to 2-trans-enoyl-CoAs (25). To verify the binding of Arid1a to their promoters, we performed ChIP-PCR with anti-Arid1a antibody in primary hepatocytes with or without *Arid1a*. Our results showed that Arid1a was present in the promoters of *Acox1* and *Cpt1a* in hepatocytes with *Arid1a*, but this occupancy was decreased in *Arid1a*-deficient cells **(Fig. 7D)**. We further explored whether Arid1a can epigenetically regulate fatty acid oxidation-related genes. Here, we detected the level of trimethylation of H3 lysine 4 (H3K4me3), an epigenetic marker associated with transcriptional activation, via ChIP-PCR with anti-H3K4me3 antibody, on *Acox1* and *Cpt1a* promoters. It turned out that H3K4me3 on these regions was significantly reduced with *Arid1a* loss **(Fig. 7E)**. Furthermore, *Arid1a* deficiency diminished the recruitment of SWI/SNF core subunit Brg1 to the promoter of *Cpt1a*, indicated by ChIP-PCR with anti-Brg1 antibody (**Fig. 7F)**. Next, we conducted ATAC-seq to assess the alterations in chromatin accessibility on the promoters of *Acox1, Cpt1a* and *Pparα*. Notably, statistical analysis on the ATAC-seq data indicated that the chromatin accessibilities on promoters of *Pparα, as well as Cpt1α*, were significantly reduced when *Arid1a* was deleted in the hepatocytes **(Fig. 7G)**. Altogether, these results indicate that Arid1a facilitates fatty acid oxidation by directly altering the epigenetic landscape of metabolism gene loci, including chromatin accessibility and local histone modification, as well as by indirectly activating downstream genes through Pparα transcription factor. *Arid1a* deficiency would result in transcriptional reduction of these FAO associated genes, thereby attenuating the corresponding metabolic functions that ends to hepatic steatosis, insulin resistance and NAFLD (**Fig. 7H**).

**Figure 7.**
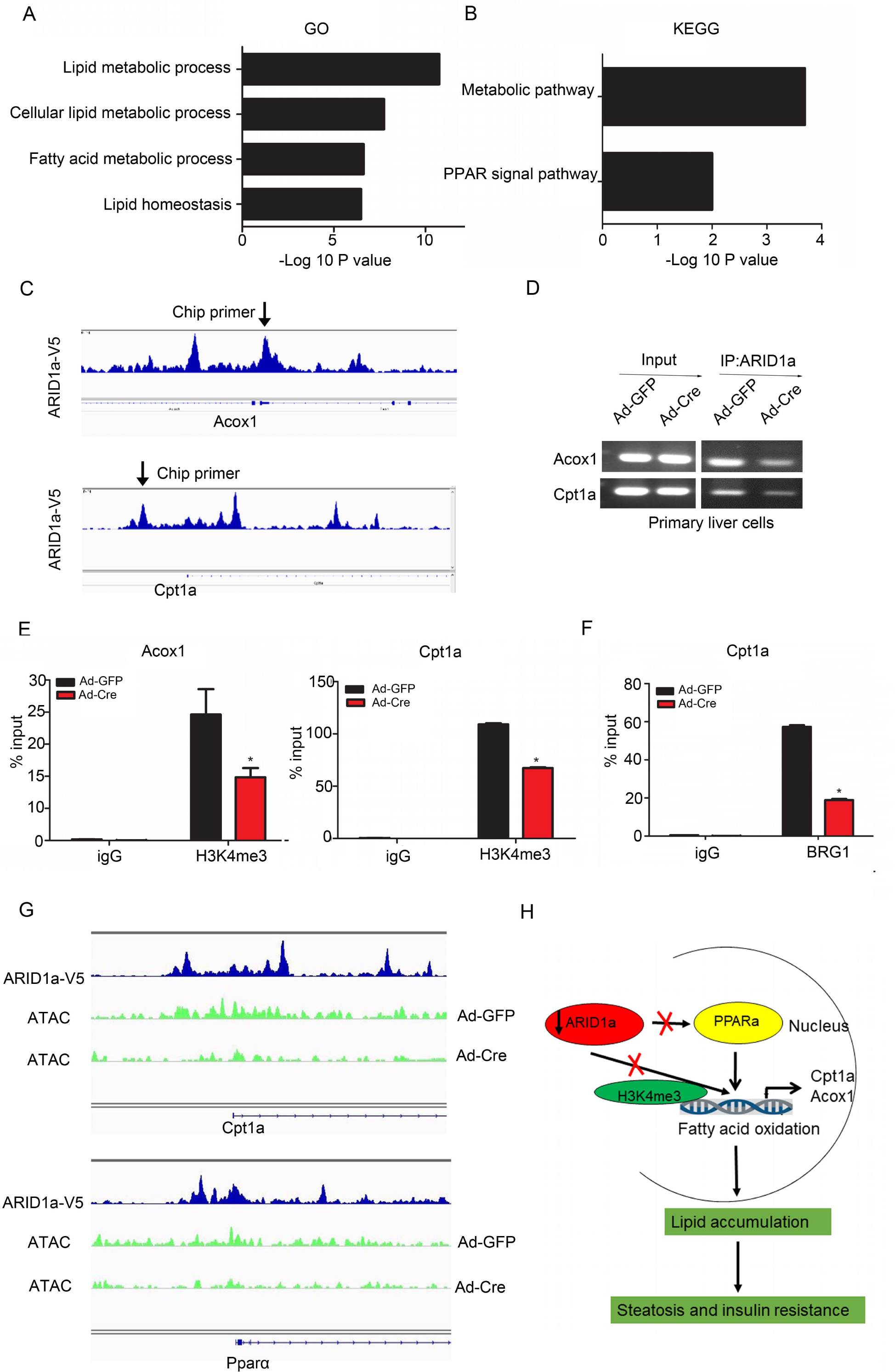
*Arid1a* deletion leads to the decreased H3K4me3 and chromatin accessibility to the metabolic genes promoter. (A-B) GO (A) and KEGG (B) pathway analysis of *Arid1a*-targeted genes identified by ChIP-Seq. (C-D) ChIP-seq (C) and ChIP-PCR (D) analysis of ARID1a-binding affinity in *Cpt1a* and *Acox1* promoter (\**P* < 0.05). (E) ChIP-qPCR analysis of H3K4me3-binding affinity in *Cpt1a* and *Acox1* promoter. Values are mean ± SD (\**P* < 0.05). (F) ChIP-qPCR analysis of BRG1-binding affinity in *Cpt1a* promoter. Values are mean ± SD (\**P* < 0.05). (G) ATAC-seq profiles show that chromatin accessibility at *Cpt1a* and *Pparα* promoters is affected by *Arid1a* depletion. (H) The proposed model for the role of Arid1a loss in hepatic steatosis and insulin resistance. Arid1a normally facilitates fatty acid oxidation by directly altering the epigenetic modification of H3K4me3 on the lipid metabolism-related genes, such as *Cpt1a, Acox1*, as well as by indirectly activating the master transcriptional factor Pparα, resulting in transcriptional activation of these FAO-associated genes, thereby maintaining lipid and glucose homeostasis. Arid1a loss disrupts the epigenetic landscape and Pparα-mediated transcription of FAO-related genes, resulting in hepatic steatosis and insulin resistance.

## Discussion

As one of the most prevalent liver diseases worldwide, NAFLD is a progressive pathological condition that promotes more severe liver and metabolic dysfunction. Lipid accumulation in the liver (steatosis) is thought to be the “first hit” of NAFLD. In this study, we identify that Arid1a is a diet-sensitive subunit of the SWI/SNF chromatin remodeling complexes in the liver, which plays an important role in increasing the susceptibility to develop hepatic steatosis and insulin resistance in mice in conditions of HFD-induced NAFLD model. At the mechanistic level, hepatic Arid1a is critical for maintaining metabolic homeostasis through Pparα-mediated fatty acid oxidation in the liver. From a clinical perspective, these data indicate that targeting ARID1a might be a promising therapeutic strategy for treating NAFLD and its related metabolic disorders.

Several lines of evidences supported a pivotal role of Arid1a in metabolic regulation. First, the transcriptional profiling and pathway analysis indicated that the genes involved in lipid metabolism, especially fatty acid metabolism were among the top down-regulated by *Arid1a* deletion. Second, mRNA levels of the genes responsible for fatty acid oxidation were significantly reduced in *Arid1a*^*-/-*^ livers and hepatocytes. Third, Arid1a downregulates *Pparα*, the major transcriptional factor for FAO (26). Consistently, Pparα in *Arid1a*^*-/-*^ hepatocytes was sufficient to reverse the lipid accumulation and insulin resistance phenotype *in vitro*. Moreover, Arid1a regulates transcription activity of key metabolic genes on the local epigenetic landscape. Specifically, ARID1a deficiency leads to reduced levels of H3K4me3 and compacted chromatin structure on promoters of key metabolic genes.

Previous studies demonstrated that SWI/SNF family member BAF60c, recruiting other subunits, including ARID1a, interacts with USF transcription factor to lipogenic gene promoters to increase lipogenesis in response to feeding/insulin treatment (8). However, in our study, Arid1a deletion leads to steatosis by downregulating expression of these genes involved in fatty acid oxidation rather than *de novo* lipogenesis pathway. Especially, the expression of Cpt1a and Acox1, two key rate-limiting enzyme in fatty acid oxidation, were largely reduced as *Arid1a* loss. It has been reported that the disorders of hepatic fatty acid oxidation lead to massive steatosis and hypertriglyceridemia in animals. Some emerging evidences have shown that hepatic FAO is also impaired in human NAFLD (19,27,28). Moreover, deficiency of PPARα or its coactivators were also shown to affect the development of NAFLD. Compared with control mice, Pparα-deficient mice developed massive steatosis, lobular inflammation, and are more susceptible to the progression of NASH due to an impairment of mitochondrial FAO, and increment of oxidative stress and inflammation (29-31). We identified Pparα expression was downregulated in hepatocytes and mice with *Arid1a* deletion, whereas overexpression of Pparα restored lipid accumulation and insulin resistance caused by *Arid1a* deficiency. Unfortunately, although by analyzing ChIP-seq data performed by others (23), Pparα contains ARID1a-binding sites on promoter, no such direct binding of ARID1a on *Ppar*αpromoter was identified in our experiment. In addition, we also did not detect any physical association between ARID1a and PPARα in 293T cells (data was not shown).

In summary, this study demonstrates that hepatocyte-specific inactivation of *Arid1a* reduced fatty acid oxidation, and aggravated diet-induced steatosis and insulin resistance in mice. Arid1a downregulates expression of the genes involved in fatty acid oxidation through Pparα and epigenetic regulation. These findings reveal a new mechanism underlying the role of Arid1a in NASH pathogenesis, and suggest a promising approach for the treatment of hepatic steatosis, fibrosis and insulin resistance by modulating Arid1a.

## Materials and methods

### Mice and diets

*Arid1a*^*LKO*^ mice were generated by mating *Arid1a*^*Flox/Flox*^ mice (kindly provided by Zhong Wang at the Cardiovascular Research Center, Massachusetts General Hospital, Harvard Medical School) with Albumin-Cre mice (from the Jackson Laboratory). *Arid1a*^*Flox/Flox*^ animals were used as controls. Six week-old *Arid1a*^*LKO*^ and *Arid1a*^*Flox/Flox*^ mice were given free access to an HFD (composed of 59% fat, 15% protein, and 26% carbohydrates based on caloric content;) or CD (protein, 20%; fat, 10%; carbohydrates, 70%;) for 12 weeks. All animal experiments were approved by the Animal Care and Use Committee of Shang Hai Jiao Tong University, and all the procedures were conducted in compliance with institutional guidelines and protocols.

### Blood parameters

The concentrations of the hepatic enzymes alanine amino transferase (ALT), aspartate amino transferase (AST), and alkaline phosphatase (ALP) were analyzed using a spectrophotometer (Chemix 180i, Sysmex Shanghai Ltd, Shanghai, China) according to manufacturer’s instructions. Insulin plasma levels were determined using mouse insulin ELISA kit (Millipore, St. Charles, MO, US). The levels of the serum inflammatory indicators were measured by ELISA (BD Bioscience, San Diego, CA, USA).

### Lipid parameters

Commercial kits were used to measure the liver triglyceride (TG), total cholesterol (TC), and nonesterified fatty acid (NEFA) contents in the liver according to the manufacturer’s instructions (290-63701 for TG, 294-65801 for TC, 294-63601 for NEFA; Wako, Tokyo, Japan). Serum lipids were analyzed using a spectrophotometer (Chemix 180i, Sysmex Shanghai Ltd, Shanghai, China) according to manufacturer’s instructions.

### Glucose and insulin tolerance tests

Glucose tolerance tests (GTT) were performed in mice that were fed either CD or HFD for 12 weeks. One week later, the same mice were used for insulin tolerance tests (ITT). For GTT, mice were fasted for 16 h. After measuring the baseline blood glucose level via a tail nick using a glucometer, 1.5g/kg glucose was administered via intraperitoneal injection, and glucose levels were measured 0,15,30,45,60,90 and 120 minutes after glucose injection. For ITT, 6-h fasted mice were injected intraperitoneally with recombinant human insulin at 1U/kg and their blood glucose concentrations were determined 0,15,30,45,60,90 and 120 minutes after insulin injection.

### Quantitative real-time PCR analysis

RNA extracted from liver or hepatocytes were subjected to reverse transcription and subsequent PCR using a real-time PCR system (ABI, Carlsbad, CA). PCR primer sequences are listed in Supplemental Table 1. Expression of the respective genes was normalized to β-Actin as an internal control.

### Primary hepatocyte isolation, cultivation and treatment

Primary hepatocytes were isolated from 6- to 8-week-old mice following the steps reported previously (32). Hepatocytes were cultured in Dulbecco’s modified Eagle’s medium contained 10% fetal bovine serum (FBS) and 1% penicillin-streptomycin in a 5% CO_2_/water-saturated incubator at 37°C. To stimulate insulin signaling, hepatocytes were stimulated with insulin 10 nM for indicated time. For lipid accumulation *in vitro*, hepatocytes were treated with 0.2 mM palmitate acid (PA, Sigma, St. Louis, MO, USA) or 0.08 mM oleic acid (OA, Sigma, St. Louis, MO, USA) for 24h.

### Plasmid Construction and Transfection

The plasmids encoding T7-ARID1a was purchased from Addgene (Cambridge, MA, USA). The plasmid expressing flag-tagged human PPARα was constructed by cloning the mouse PPARα cDNA into the pcDNA3.1 vector. Transfection assays for plasmids were performed using Lipofectamine 2000 (Invitrogen; Carlsbad, CA, US) according to the manufacturer’s protocol.

### Histological analysis

Sirius red (Sigma, St. Louis, MO, USA) staining was performed using paraffin-prepared liver sections to determine the fibrosis of the tissues. Liver sections were embedded in paraffin and stained with hematoxylin and eosin (H&E) to visualize adipocytes and inflammatory cells in the tissues. Frozen liver sections of 10mm were fixed in formalin and then rinsed with 60% isopropanol. After staining with freshly prepared Oil-red O solution (Sigma, St. Louis, MO, USA) for 10min, the sections were rinsed again with 60% isopropanol. Sections or cells were analyzed by inverted microscope (Olympus, Tokyo, Japan).

### Western blotting

After extracted from liver samples or cultured cells, each protein sample (50 mg) was subjected to SDS/PAGE and transferred to NC membrane (Millipore, Bedford, MA, USA). Then the corresponding primary and secondary antibodies were incubated to visualize the protein. Antibodies used in western blot analysis included anti-ARID1a (12354, CST, Billerica, MA, USA), anti-phospho-AKTS473 (9271, CST, Billerica, MA, USA), anti-phospho-GSK3β (9315, CST, Billerica, MA, USA), anti-AKT (9272, CST, Billerica, MA, USA), anti-GSK3β (9323, CST, Billerica, MA, USA), anti-PPARα (15540-I-AP, ProteinTech, Chicago, IL, USA), anti-β-actin (2381, (Sigma, St. Louis, MO, USA)).

### ChIP

Chromatin immunoprecipitation (ChIP) assay was performed according to the protocol developed by Upstate Biotechnology as described (11). Briefly, chromatin lysates were prepared from hepatocytes following crosslinking with 1% formaldehyde. The samples were precleared with Protein-G agarose beads and immunoprecipitated using antibodies against ARID1a (sc32761-x, Santa Cruz, California, USA), H3K4me3 (Abcam, 8580, Cambridge, UK),BRG1 (sc17796, Santa Cruz, Cruz, California, USA) or control IgG in the presence of BSA and salmon sperm DNA. Beads were extensively washed before reverse crosslinking. DNA was purified using a PCR Purification Kit (Tiangen,Beijing,China) and subsequently analyzed by qPCR using primers located on the Acox1 and Cpt1a promoter.

### RNA-Seq

Total RNA-seq was performed on three independent biological replicates per condition. Samples from different conditions were processed together to prevent batch effects. For each sample, 10 million cells were used to extract total RNA and produce RNA-seq libraries. Raw sequencing reads were trimmed by Trimmomatic (v0.36) to remove adapters and low quality sequences. The filtered reads were aligned to the GRCm38 UCSC annotated transcripts via Tophat (v2.1.0)56. Transcripts were then assembled, counted and normalized with the Cufflinks suit (v2.2.1)57. Differentiated expressed genes were analyzed by Cuffdiff in the Cufflinks suit, using p value <0.05 and fold change >1.5 as cutoff. Heatmaps were generated by using R package pheatmap (1.0.10).

Gene ontology and KEGG pathway enrichment analyses were performed using DAVID (v6.8) with the differentially expressed gene lists. Gene set enrichment analysis (GSEA v3.0) was performed using the normalized expression values. The raw sequencing data of RNA-seq has been deposited to NCBI GEO database with the accession number (GSE….)

### ATAC-seq

Cells were harvested and subjected for ATAC-seq analysis as previously described (33). Briefly, 50, 000 cells were washed with cold PBS and cell membrane was lysed in lysis buffer (10 mM Tris-HCl, pH 7.4, 10 mM NaCl, 3 mM MgCl_2_, 0.1% Nonidet P-40). After centrifugation, cell pellets were resuspended and incubated in transposition mix containing Tn5 transposase (Illumina, California, USA) at 37 °C for 30 minutes. Purified DNA was then ligated with adapters and amplified 11 total cycles. Libraries were purified with AMPure beads (Beckman Coulter Inc, Brae, CA, USA) to remove contaminating primer dimers. Libraries were then sequenced on Illumina X-Ten system with 150 bp paired-end sequencing strategy.

For ATAC-seq data analysis, Illumina adapters and low-quality reads were trimmed by Trimmomatic (v0.36) (Trimmomatic: a flexible trimmer for illumina sequence data). Afterwards, all reads for each sample were combined and aligned to mouse reference build GRCm38 using bwa (v0.7.11) with default settings. More than 80 million pair-ended sequencing reads were obtained from each library of the MEF cells and 40 to 65 million reads were obtained from the libraries of the primary mouse liver cells. Low quality reads (mapping quality < 20) and reads mapping to mitochondrial DNA were removed using samtools (v1.6). Duplicates were excluded using Picard Tools (v1.4.5). Between 49 and 61 million high-quality reads per sample that mapped to genomic DNA were obtained for MEF cells. Between 11 and 25 million high-quality reads per sample that mapped to genomic DNA were obtained for mouse primary liver cells. All mapped reads were offset by +4bp for the +strand and -5bp for the –strand (Transposition of native chromatin for fast and sensitive epigenomic profiling of open chromatin, DNA-binding proteins and nucleosome position) using bedtools (v2.25.0). Libraries of the same condition were merged for subsequent analyses. Peaks were called for each sample using MACS2 (Model-based analysis of ChIP-Seq (MACS)) (v2.1.1.20160309) using parameter “-–nomodel –shift 100 –extsize 200”. The peaks with p<0.01 for each group in the same cell line were merged for searching differential peaks. Differential peak calling, as well as peak annotation and motif analysis were performed using HOMER (v4.9.1) with default settings. Fold change larger than 2 was set as the cutoff for differential peak identification. Sequencing signals were generated by transforming the mapping files into bigwig tracks, which was visualized in the Integrated Genomic Viewer (IGV). Signal heat maps centered around peaks were generated using deep Tools (v 2.5.3). All genomic datasets were deposited in GEO datasets with the accession number (GSE……).

### Statistics

All statistical analyses were performed using Graphpad prism software (version 19.0), and all data are expressed as the mean ± standard deviation. For data with a normal distribution and homogeneity of variance, two-tailed Student t tests were used to evaluate significant differences between two groups, *P* values less than 0.05 were considered significant. * *P* < 0.05; ** *P* < 0.01; *** *P* < 0.001.

## Supplementary information

Supplemental Information includes four figures and one table can be found with this article online at

## Acknowledgments

We sincerely thank Zhong Wang professor at the Cardiovascular Research Center of Harvard Medical School, for providing *Arid1a*^*F/F*^ mice in this study. This work was supported by National Natural Science Foundation of China (81672772 and 81472621), China National Science and Technology Major Project for Prevention and Treatment of Infectious Diseases (grant no. 2017ZX10203207) and National Program on Key Research Project of China (grant no. 2016YFC0902701)

## Abbreviations

NAFLD: non-alcoholic fatty liver disease
NASH: non-alcoholic steatohepatitis
FAO: fatty acid oxidation
FAs: fatty acids
FD: high fat diet
CD: chow diet
GTT: glucose tolerance test
ITT: insulin tolerance test
LKO: liver-specific knockout
HE: hematoxylin and eosin
TG: triglyceride
TCHO: total cholesterol
NEFA: non-esterified fatty acid
ALT: alanine amino transferase
AST: aspartate amino transferase
ALP: alkaline phosphatase
KEGG: Kyoto Encyclopedia of Genes and Genomes
GO: gene ontology
OA: oleic acid
PA: palmitic acid.

## References

1. Loomba, R., and Sanyal, A. J. (2013) The global NAFLD epidemic. Nature reviews. Gastroenterology & hepatology 10, 686–690

2. Demir, M., Lang, S., and Steffen, H. M. (2015) Nonalcoholic fatty liver disease - current status and future directions. Journal of digestive diseases 16, 541–557

3. Samuel, V. T., and Shulman, G. I. (2018) Nonalcoholic Fatty Liver Disease as a Nexus of Metabolic and Hepatic Diseases. Cell metabolism 27, 22–41

4. Fritz, I. B., and Yue, K. T. (1963) Long-Chain Carnitine Acyltransferase and the Role of Acylcarnitine Derivatives in the Catalytic Increase of Fatty Acid Oxidation Induced by Carnitine. Journal of lipid research 4, 279–288

5. Muchardt, C., and Yaniv, M. (1999) ATP-dependent chromatin remodelling: SWI/SNF and Co. are on the job. Journal of molecular biology 293, 187–198

6. Wilson, B. G., and Roberts, C. W. (2011) SWI/SNF nucleosome remodellers and cancer. Nature reviews. Cancer 11, 481–492

7. Gresh, L., Bourachot, B., Reimann, A., Guigas, B., Fiette, L., Garbay, S., Muchardt, C., Hue, L., Pontoglio, M., Yaniv, M., and Klochendler-Yeivin, A. (2005) The SWI/SNF chromatin-remodeling complex subunit SNF5 is essential for hepatocyte differentiation. The EMBO journal 24, 3313–3324

8. Wang, Y., Wong, R. H., Tang, T., Hudak, C. S., Yang, D., Duncan, R. E., and Sul, H. S. (2013) Phosphorylation and recruitment of BAF60c in chromatin remodeling for lipogenesis in response to insulin. Molecular cell 49, 283–297

9. Meng, Z. X., Li, S., Wang, L., Ko, H. J., Lee, Y., Jung, D. Y., Okutsu, M., Yan, Z., Kim, J. K., and Lin, J. D. (2013) Baf60c drives glycolytic metabolism in the muscle and improves systemic glucose homeostasis through Deptor-mediated Akt activation. Nature medicine 19, 640–645

10. Meng, Z. X., Wang, L., Chang, L., Sun, J., Bao, J., Li, Y., Chen, Y. E., and Lin, J. D. (2015) A Diet-Sensitive BAF60a-Mediated Pathway Links Hepatic Bile Acid Metabolism to Cholesterol Absorption and Atherosclerosis. Cell reports 13, 1658–1669

11. Li, S., Liu, C., Li, N., Hao, T., Han, T., Hill, D. E., Vidal, M., and Lin, J. D. (2008) Genome-wide coactivation analysis of PGC-1alpha identifies BAF60a as a regulator of hepatic lipid metabolism. Cell metabolism 8, 105–117

12. Dallas, P. B., Cheney, I. W., Liao, D. W., Bowrin, V., Byam, W., Pacchione, S., Kobayashi, R., Yaciuk, P., and Moran, E. (1998) p300/CREB binding protein-related protein p270 is a component of mammalian SWI/SNF complexes. Molecular and cellular biology 18, 3596–3603

13. Dallas, P. B., Pacchione, S., Wilsker, D., Bowrin, V., Kobayashi, R., and Moran, E. (2000) The human SWI-SNF complex protein p270 is an ARID family member with non-sequence-specific DNA binding activity. Molecular and cellular biology 20, 3137–3146

14. Martens, J. A., and Winston, F. (2003) Recent advances in understanding chromatin remodeling by Swi/Snf complexes. Current opinion in genetics & development 13, 136–142

15. Nagl, N. G., Jr., Zweitzig, D. R., Thimmapaya, B., Beck, G. R., Jr., and Moran, E. (2006) The c-myc gene is a direct target of mammalian SWI/SNF-related complexes during differentiation-associated cell cycle arrest. Cancer research 66, 1289–1293

16. Trotter, K. W., and Archer, T. K. (2004) Reconstitution of glucocorticoid receptor-dependent transcription in vivo. Molecular and cellular biology 24, 3347–3358

17. Trotter, K. W., Fan, H. Y., Ivey, M. L., Kingston, R. E., and Archer, T. K. (2008) The HSA domain of BRG1 mediates critical interactions required for glucocorticoid receptor-dependent transcriptional activation in vivo. Molecular and cellular biology 28, 1413–1426

18. Fang, J. Z., Li, C., Liu, X. Y., Hu, T. T., Fan, Z. S., and Han, Z. G. (2015) Hepatocyte-Specific Arid1a Deficiency Initiates Mouse Steatohepatitis and Hepatocellular Carcinoma. PloS one 10, e0143042

19. Samuel, V. T., and Shulman, G. I. (2012) Mechanisms for insulin resistance: common threads and missing links. Cell 148, 852–871

20. Kleiner, D. E., Brunt, E. M., Van Natta, M., Behling, C., Contos, M. J., Cummings, O. W., Ferrell, L. D., Liu, Y. C., Torbenson, M. S., Unalp-Arida, A., Yeh, M., McCullough, A. J., Sanyal, A. J., and Nonalcoholic Steatohepatitis Clinical Research, N. (2005) Design and validation of a histological scoring system for nonalcoholic fatty liver disease. Hepatology 41, 1313–1321

21. Fabbrini, E., Sullivan, S., and Klein, S. (2010) Obesity and nonalcoholic fatty liver disease: biochemical, metabolic, and clinical implications. Hepatology 51, 679–689

22. Leone, T. C., Weinheimer, C. J., and Kelly, D. P. (1999) A critical role for the peroxisome proliferator-activated receptor alpha (PPARalpha) in the cellular fasting response: the PPARalpha-null mouse as a model of fatty acid oxidation disorders. Proceedings of the National Academy of Sciences of the United States of America 96, 7473–7478

23. Sun, X., Chuang, J. C., Kanchwala, M., Wu, L., Celen, C., Li, L., Liang, H., Zhang, S., Maples, T., Nguyen, L. H., Wang, S. C., Signer, R. A., Sorouri, M., Nassour, I., Liu, X., Xu, J., Wu, M., Zhao, Y., Kuo, Y. C., Wang, Z., Xing, C., and Zhu, H. (2016) Suppression of the SWI/SNF Component Arid1a Promotes Mammalian Regeneration. Cell stem cell 18, 456–466

24. Bergman, A. J., Donckerwolcke, R. A., Duran, M., Smeitink, J. A., Mousson, B., Vianey-Saban, C., and Poll-The, B. T. (1994) Rate-dependent distal renal tubular acidosis and carnitine palmitoyltransferase I deficiency. Pediatric research 36, 582–588

25. Zeng, J., and Li, D. (2004) Expression and purification of his-tagged rat peroxisomal acyl-CoA oxidase I wild-type and E421 mutant proteins. Protein expression and purification 38, 153–160

26. McGarry, J. D., and Foster, D. W. (1980) Regulation of hepatic fatty acid oxidation and ketone body production. Annual review of biochemistry 49, 395–420

27. Schmid, A. I., Szendroedi, J., Chmelik, M., Krssak, M., Moser, E., and Roden, M. (2011) Liver ATP synthesis is lower and relates to insulin sensitivity in patients with type 2 diabetes. Diabetes care 34, 448–453

28. Cortez-Pinto, H., Chatham, J., Chacko, V. P., Arnold, C., Rashid, A., and Diehl, A. M. (1999) Alterations in liver ATP homeostasis in human nonalcoholic steatohepatitis: a pilot study. Jama 282, 1659–1664

29. Ip, E., Farrell, G. C., Robertson, G., Hall, P., Kirsch, R., and Leclercq, I. (2003) Central role of PPARalpha-dependent hepatic lipid turnover in dietary steatohepatitis in mice. Hepatology 38, 123–132

30. Abdelmegeed, M. A., Yoo, S. H., Henderson, L. E., Gonzalez, F. J., Woodcroft, K. J., and Song, B. J. (2011) PPARalpha expression protects male mice from high fat-induced nonalcoholic fatty liver. The Journal of nutrition 141, 603–610

31. Su, Q., Baker, C., Christian, P., Naples, M., Tong, X., Zhang, K., Santha, M., and Adeli, K. (2014) Hepatic mitochondrial and ER stress induced by defective PPARalpha signaling in the pathogenesis of hepatic steatosis. American journal of physiology. Endocrinology and metabolism 306, E1264–1273

32. Kegel, V., Deharde, D., Pfeiffer, E., Zeilinger, K., Seehofer, D., and Damm, G. (2016) Protocol for Isolation of Primary Human Hepatocytes and Corresponding Major Populations of Non-parenchymal Liver Cells. Journal of visualized experiments : JoVE, e53069

33. Buenrostro, J. D., Wu, B., Chang, H. Y., and Greenleaf, W. J. (2015) ATAC-seq: A Method for Assaying Chromatin Accessibility Genome-Wide. Current protocols in molecular biology 109, 21 29 21–29

