## Supplementary figures and for "*Arid1a* protects against hepatic steatosis and insulin resistance via PPARα-mediated fatty acid oxidation"

**Supplementary Figure 1.** ***Arid1a*** $^{LKO}$ **mice does not exhibit insulin resistance phenotypes on normal chow diet.**

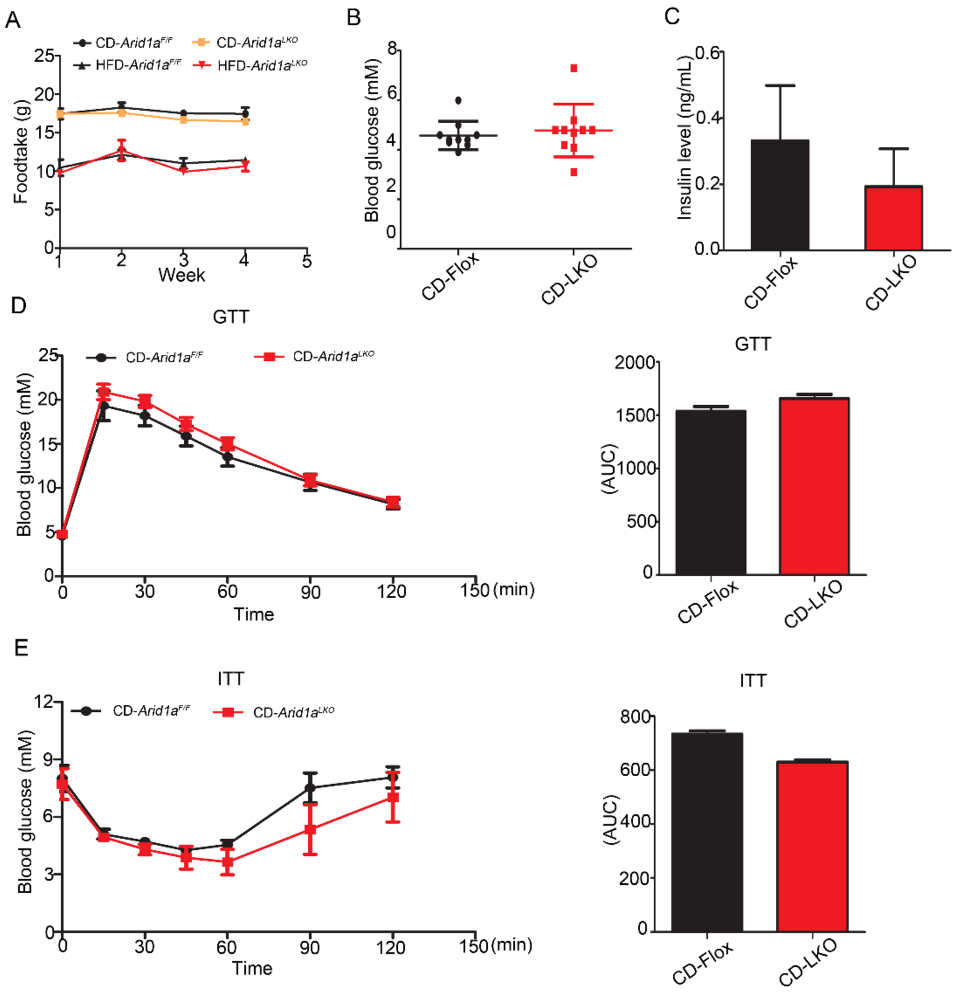


**Supplementary Fig 1.** ***Arid1a*** $^{LKO}$ **mice does not exhibit insulin resistance phenotypes on normal chow diet.**

(A) Food intake (n=10).

(B) Fasting blood glucose levels of mice on CD (n=10).

(C) ELISA-determined fasting insulin levels of mice on CD (n=8).

(D) Left, measurement of plasma glucose during glucose tolerance test of mice. Right, the calculated AUCs (area under cureves) in mice on CD (n=10).

(E) Left, measurement of plasma glucose during insulin tolerance test of mice. Right, the calculated AUCs (area under cureves) in mice on CD (n=6-7).

Supplementary Figure 2. *Arid1a* deficiency leads to insulin resistance.

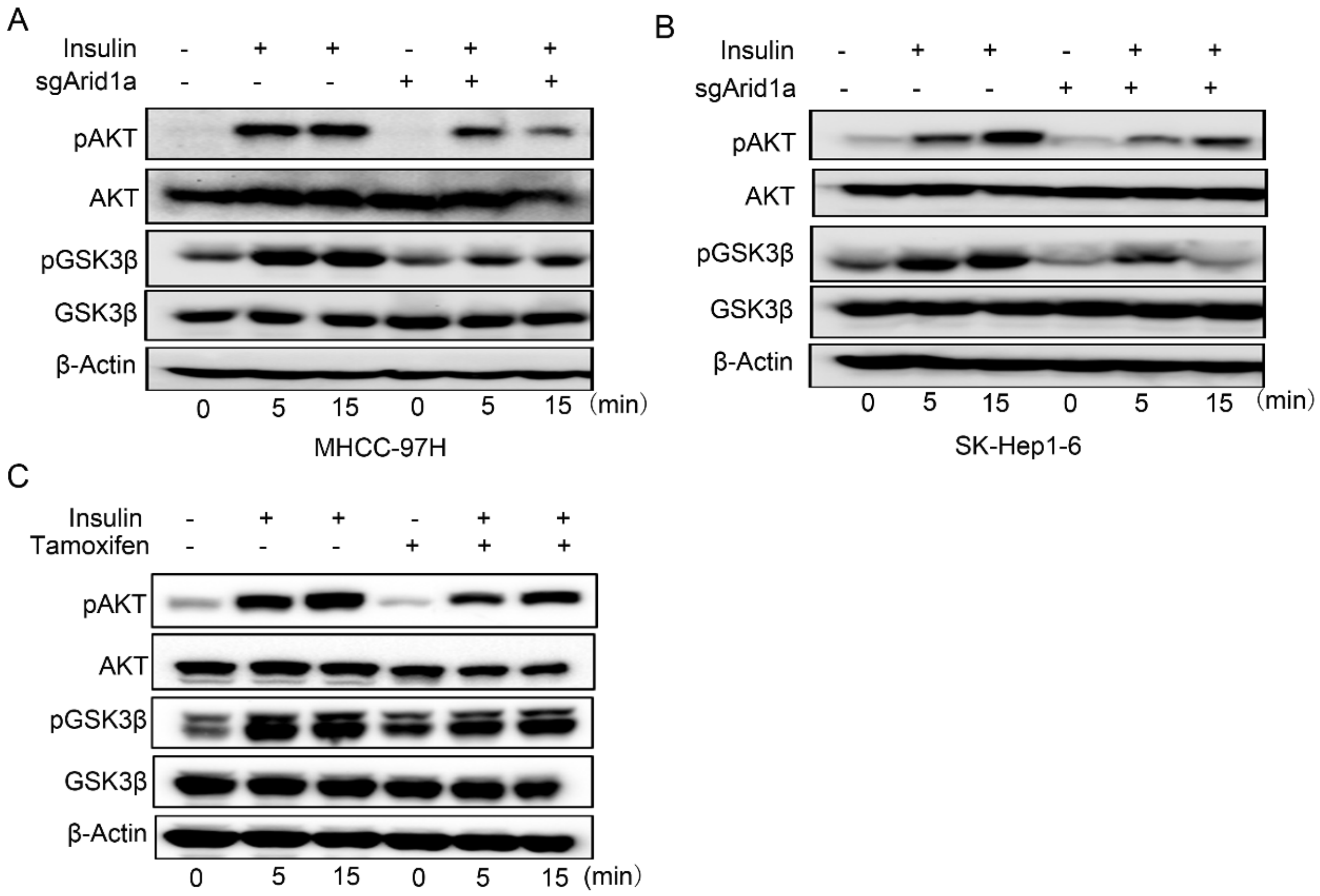


**Supplementary Figure 2. *Arid1a* deficiency leads to insulin resistance.**

(A-C) MHCC-97H (A), SK-Hep1-6 (B) and primary hepatocytes isolated from tamoxifen-induced ***Arid1a*** $^{LKO}$ mice (C) were stimulated with insulin (10 nM) for the indicated times, and phosphorylation of AKT and GSK were determined.

Supplementary Figure 3. *Arid1a* deficiency leads to FAO defieciency.

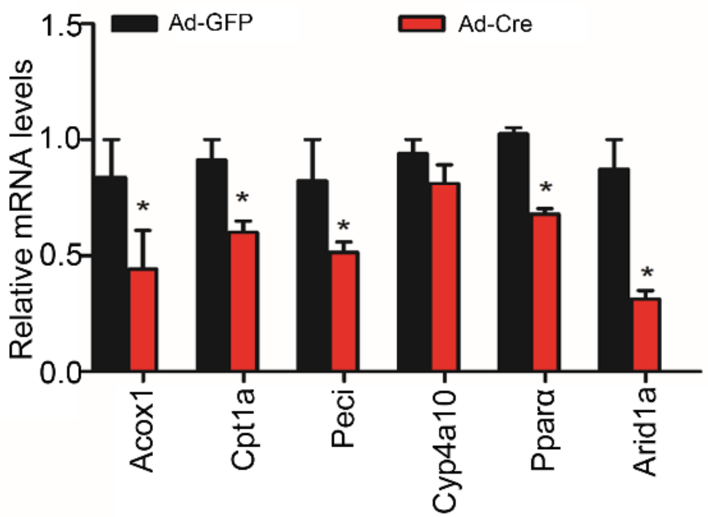


**Supplementary Figure 3. *Arid1a* deficiency leads to FAO defieciency.**

Real-time analysis of genes involved in fatty acid oxidation in hepatocytes immortalizated with SV40 (*$p<0.05$). Values are mean ± SD.

.

**Supplementary Figure 4. *Arid1a* deletion downregulates Irs1 expression.**

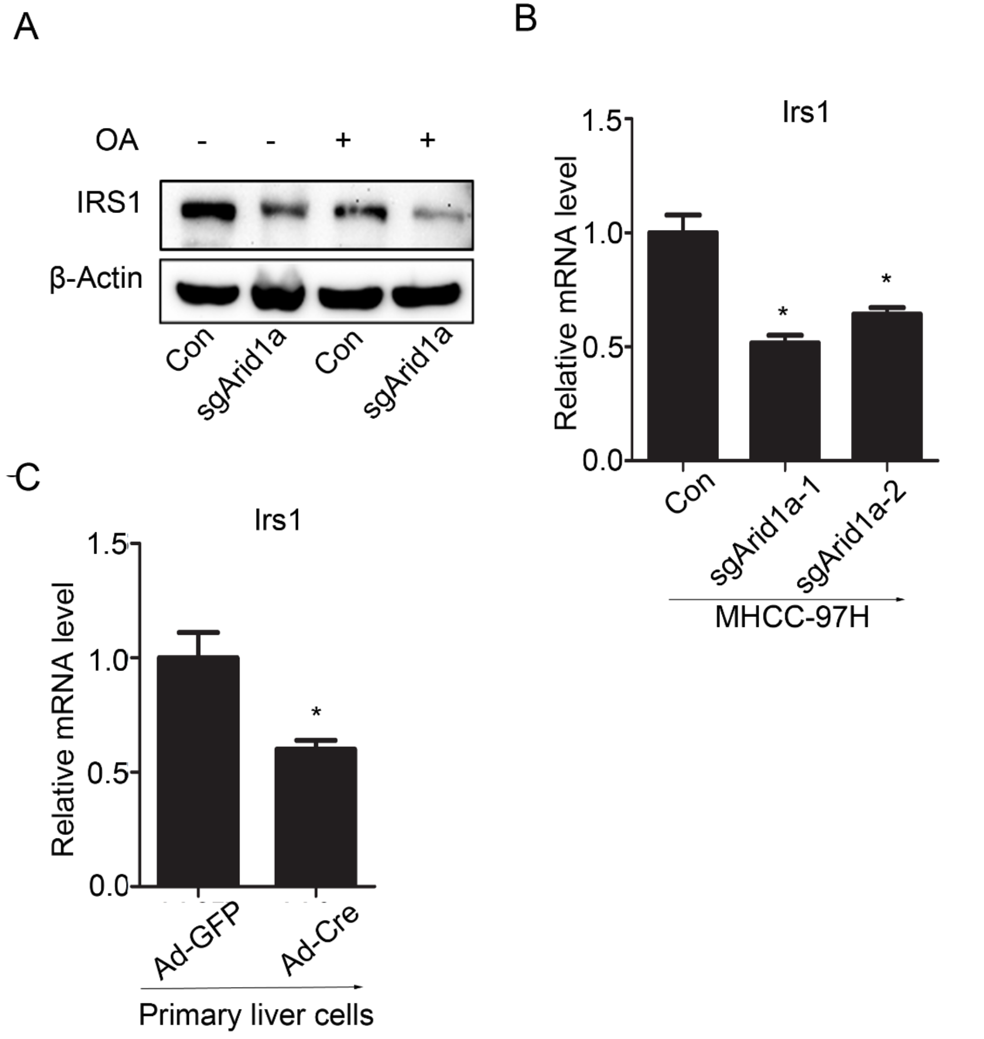


**Supplementary Figure 4. *Arid1a* deletion downregulates Irs1 expression.**

(A) Protein expression level of Irs1 in MHCC-97H.

(B-C) mRNA expression levels of Irs1 in MHCC-97H (B) and hepatocytes (C) (*$p<0.05$). Values are mean ± SD.

**Table 1. List of primer sequences used in real-time PCR**

| Gene name | sequence (5'to3') |
|---|---|
| Acox1-F | CCTGATTCAGCAAGGTAGGG |
| Acox1-R | TCGCAGACCCTGAAGAAATC |
| Cpt1a-F | AGTGGCCTCACAGACTCCAG |
| Cpt1a-R | GCCCATGTTGTACAGCTTCC |
| Hmgcs2-F | ATACCACCAACGCCTGTTATG |
| Hmgcs2-R | CAATGTCACCACAGACCACCA |
| Ppara-F | TTTCGGCGAACTATTCGGCTG |
| Ppara-R | GGCATTTGTTCCGGTTCTTCTT |
| Cyp4a10-F | AAGGGTCAAACACCTCTGGA |
| Cyp4a10-R | GATGGACGCTCTTTACCCAA |
| Pparr-F | GCTGTTATGGGTGAAACTCT |
| Pparr-R | TGGCATCTCTGTGTCAACCA |
| Cox7a1-F | GTCTCCCAGGCTCTGGTCCG |
| Cox7a1-R | CTGTACAGGACGTTGTCCATTC |
| Peci-F | CGAGTTGGCTGAATGGAGTA |
| Peci-R | CCAGCTGTGGGAATCTCTGT |
| Pgc1a-F | TCCTCCTCATAAAGCCAACC |
| Pgc1a-R | GCCTTGGGTACCAGAACACT |
| Mcad-F | CCAGAGAGGAGATTATCCCCG |
| Mcad-R | TACACCCATACGCCAACTCTT |
| Mttp-F | GACCACCCTGGATCTCCATA |
| Mttp-R | AGCGTGGTGAAAGGGCTTAT |
| Mcp1-F | TTTTTGTCACCAAGCTCAAGAGA |
| Mcp1-R | ATTTGGTTCCGATCCAGGTT |
| Tnfa-F | CATCTTCTCAAAATTCGAGTGACAA |
| Tnfa-R | TGGGAGTAGACAAGGTACAACCC |
| Cd11b-F | ATCAACACAACCAGAGTGGATTC |
| Cd11b-R | GTTCCTCAAGATGACTGCAGAAG |
| Arg1-F | AGACCACAGTCTGGCAGTTG |
| Arg1-R | CCACCCAAATGACACATAGG |
| IL6-F | TTCCATCCAGTTGCCTTCTTGG |
| IL6-R | TTCTCATTTCCACGATTTCCCAG |
| Srebp1c-F | GTTACTCGAGCCTGCCTTCAGG |
| Srebp1c-R | CAAGCTTTGGACCTGGGTGTG |
| Acc1-F | GGACAGACTGATCGCAGAGAAAG |
| Acc1-R | TGGAGAGCCCCACACACA |
| Fas-F | GCTGCGGAAACTTCAGGAAAT |
| Fas-R | AGAGACGTGTCACTCCTGGACTT |
| Pepck-F | GTGGGAGTGACACCTCACAGC |
| Pepck-R | AGGACAGGGCTGGCCGGGACG |
| G6p-F | ATGAACATTCTCCATGACTTTGGG |
| G6p-R | GACAGGGAACTGCTTTATTATAGG |
